# Mosquito Viromes in England and Wales Reveal Hidden Arbovirus Signals and Limited Ecological Structuring

**DOI:** 10.1101/2025.11.11.687861

**Authors:** Jack Pilgrim, Emma Widlake, Roksana Wilson, Alexander G C Vaux, Jolyon M Medlock, Alistair C Darby, Matthew Baylis, Marcus SC Blagrove

## Abstract

Outbreaks of mosquito-borne viruses are increasing in temperate regions, with West Nile and Usutu viruses now established in wide regions across Europe, and both detected in the UK. Current surveillance strategies focus on targeted approaches which are well suited for monitoring established threats but limited in their ability to detect recently described or neglected viruses. High throughput sequencing (HTS) provides an unbiased alternative, allowing simultaneous identification of well-recognised and overlooked arboviruses, alongside insect-specific viruses (ISVs) that may modulate vector competence of the insects transmitting these pathogens.

This study presents the first comprehensive virome survey of *Culex* mosquitoes in the UK, analysing populations collected from 93 sites across England and Wales through HTS and a systematic virus discovery pipeline. Across these sites, 41 distinct viral taxa were identified, including 11 novel species. Most viruses were rare or confined to a few sites, with only three detected in more than one third of sites, suggesting the absence of a broad conserved virome across populations. Within this diversity, three arbovirus-related lineages were detected: Hedwig virus (*Peribunyaviridae*), Umatilla virus (*Sedoreoviridae*), and Atherstone virus (*Peribunyaviridae*), the former two representing the first detections in the UK. These putative arboviruses were embedded in viral communities that showed minimal structuring by coarse land type but a modest decline in richness with latitude across rural sites, consistent with diversity gradients observed in other microbial systems.

Together, these findings provide the first national-scale baseline of *Culex* mosquito-associated viral diversity in the UK, and demonstrate the value of metagenomic approaches in arbovirus preparedness.

## Introduction

Arboviral activity in Europe has intensified in recent years, with previously sporadic detections giving way to sustained transmission in some regions [1] and novel viruses appearing in areas where they were historically absent [2]. Usutu virus (USUV) and West Nile virus (WNV) are now established across parts of central and southern Europe [3–5], with USUV causing repeated epizootics in wild birds [6] and WNV showing seasonal transmission in countries such as Italy, Greece, and Spain [7–10]. In the UK, USUV became the first enzootic mosquito-borne virus following its detection in birds and mosquitoes in 2020 [11], and in 2023, WNV was detected in mosquitoes for the first time [12], reflecting the country’s growing alignment with broader European arbovirus trends. While USUV and WNV are currently viewed as the primary mosquito-borne threats, other arboviruses, including alphaviruses (e.g. Sindbis virus [13] and chikungunya virus [14]) and orthobunyaviruses [15–17], such as Tahyna virus, have also been reported in European mosquito populations. This highlights the wide range of arboviruses circulating across the continent, which may pose future emergence risks for the UK [18,19].

Despite these detections, the UK has yet to conduct a large-scale survey of mosquito-associated viral diversity. In common with most regions, current surveillance remains focused on a small number of established threats, primarily through targeted PCR or vertebrate serology [20]. To address this gap, high-throughput sequencing (HTS) offers a powerful and unbiased alternative, enabling simultaneous detection of recognised arboviruses, highly divergent taxa, and viruses with no prior association to mosquitoes [21]. Recent metagenomic investigations from Europe [22–24], Asia [25–27], and the Americas [28,29] have confirmed that mosquito populations harbour unexpectedly rich viral communities, including insect-specific viruses (ISVs) and novel lineages of uncertain host range or pathogenic potential.

Among these detections, ISVs have received growing attention due to their demonstrated ability to modulate arbovirus replication and transmission [30,31]. For example, several insect-specific flaviviruses have been shown to reduce dissemination or replication of Zika, West Nile, and dengue viruses in both *Aedes* and *Culex* mosquitoes [32,33]. The proposed mechanisms include superinfection exclusion [34], in which closely related viruses compete for similar replication niches and cellular factors, or a broader immune priming effect through activation of host antiviral pathways [31]. Despite uncertainty about their role in wild populations, ISVs are increasingly investigated as candidates for biocontrol strategies [35,36].

We previously used metagenomic sequencing to investigate mosquito viromes at two UK zoos, identifying 26 viruses, including the first report of two novel orthobunyaviruses with putative arboviral potential [37]. However, the restricted geographic scope of that study limited inferences about viral prevalence, and ecological drivers of diversity across the UK.

Here, we build on that work by conducting the first comprehensive *Culex* spp. virome survey across England and Wales, analysing mosquitoes collected from 93 sites. The objectives of this study were to (i) characterise the diversity and phylogenetic relationships of viruses associated with native *Culex* populations, (ii) examine spatial and ecological patterns in virome composition, (iii) identify candidate ISVs that may influence vector competence, and (iv) detect viruses of possible relevance to animal or public health. In doing so, we provide the first national-scale assessment of mosquito-associated viral diversity in the UK, highlighting the diversity of viruses present in mosquito populations across the region and informing future surveillance strategies.

## Methods

### Mosquito collections and pooling

Adult mosquitoes were obtained during July 2023 as part of a *Culex* trapping project covering 200 sites across England and Wales (see [38]). Trapping employed BG-PRO® traps (Biogents AG, Regensburg, Germany) baited with BG-Lure® and BG-CO₂ Generators, together with BG-GAT® gravid traps, which were operated for 72 h at each site. Mosquitoes were stored at –80 °C until processing.

Specimens belonging to the *Culex pipiens* complex and *Culex torrentium* were identified morphologically and confirmed by PCR as previously described [38]. Between 1 and 10 individuals per site were combined to form a pool, depending on site yields (See supplemental data for pooling and collection information). Where >10 mosquitoes were collected from a site, multiple replicates were prepared. Whole mosquitoes were homogenised using a bead beater (5 m/s, 40 s) with 2 mm silica beads in 100 µl Proteinase K buffer (Life Sciences). Of this, 50 µl was reserved for species identification, and 50 µl was retained for pooling. Pooled volumes were adjusted to 500 µl with 1× PBS where required (if under 10 individuals). Only females were included in virome sequencing.

In total, 151 pools representing 93 sites were generated for sequencing (Totalling 948 individuals). A PBS-only sample was included as a negative control. For the positive control, a *Culex pipiens molestus* female was fed on a blood meal containing Usutu virus at a final concentration of 4.0 × 10⁷ pfu/ml, corresponding to an estimated dose of ∼4.0 × 10⁴ pfu per mosquito (assuming ingestion of ∼1 µl of blood).

### Nucleic acid extraction and viral RNA enrichment

Pooled homogenates were centrifuged at 16,000 × g for 5 min at 4 °C, and 300 µl of clarified supernatant was filtered through a 0.45 µm sterile spin filter (Corning Costar Spin). If clogging occurred, the remaining material was transferred to a fresh spin column until all supernatant was processed.

Filtered homogenates were treated with 2 units TURBO DNase (Thermo Fisher Scientific) to remove host and bacterial DNA. RNA was purified using RNAClean xp beads (Beckman Coulter) according to the manufacturer’s instructions. Ribosomal RNA was depleted using the NEBNext rRNA Depletion Kit (New England Biolabs), supplemented with custom probes targeting conserved *Culex* rRNA regions. Depletion followed the manufacturer’s protocol with the addition of the mosquito-specific probes.

RNA quality and fragment size distribution were assessed using an Agilent 5300 Fragment Analyzer, and concentrations were determined using a Qubit™ RNA HS (High Sensitivity) Assay Kit (Thermo Fisher Scientific). Reverse transcription and sequence-independent single primer amplification (SISPA) was conducted to enrich viral RNA following the modified protocol described in Pilgrim et al., [37].

Libraries were prepared using the NEBNext Ultra II FS DNA Library Prep Kit for Illumina (New England Biolabs), incorporating fragmentation, end repair, adaptor ligation, and indexing. Clean-up was performed with AMPure XP beads. Libraries were quantified with a Qubit™ 1X dsDNA High Sensitivity assay kit and fragment distributions verified with an Agilent 5300 Fragment Analyzer prior to sequencing.

### Illumina sequencing

All libraries were sequenced on two lanes of the Illumina NovaSeq X Plus platform using 25B chemistry with 150 bp paired-end reads, generating 3.332 billion reads.

### Read processing and assembly

Illumina adapter and SISPA primer sequences were trimmed from raw FASTQ files using Cutadapt version 4.5 [39]. Reads were further trimmed to remove low quality bases with a minimum window quality score of 20. Reads shorter than 15 bp were then removed and sequencing quality was assessed with FastQC v0.12.1 [40]. De novo assembly was carried out using MEGAHIT v1.2.9 [41] with default parameters, and only contigs longer than 1,000 nucleotides were retained for further analysis.

### Initial viral signal detection

Putative viral sequences were identified using a combination of homology- and signature-based approaches. First, contigs were compared against the Virus-Host DB virus [42] protein database using BLAST+ v2.15.0 [42], with an e-value cut-off of 1 × 10⁻⁵ and a minimum query coverage per high-scoring segment of 30%. In parallel, VirSorter2 v2.2.3 [43] was run with default parameters to detect RNA viruses. Any contig identified as viral by either method was carried forward to subsequent steps.

### Post-assembly re-construction and viral gene detection

To improve contiguity, candidate viral contigs were processed with Contig Overlap Based Re-Assembly (COBRA) [44]. Protein-coding genes were predicted from these extended contigs using Prodigal v2.6.3 [45] with the “meta” mode, and the resulting protein sequences were screened against RVDB-prot v29.0 [46] using HMMsearch (HMMER v3.3.2 [47]). Contigs containing proteins with significant similarity to viral families (e-value ≤ 1 × 10⁻⁵) were retained.

### Completeness estimation and filtering

Viral contigs were evaluated for genome completeness using ViralQC [48]. Those with an estimated completeness of at least 50% were retained. Contigs not scored by ViralQC were assessed using a rescue pipeline in which predicted proteins were queried against a custom ICTV-derived NR protein database with MMseqs2 v14.7e284 [49], and taxonomy was assigned using a lowest common ancestor approach. Completeness was estimated from MMseqs2 assignments, and the same ≥50% threshold was applied. Results from both approaches were integrated to yield a high-confidence viral contig set.

### Dereplication and genome filtering

To reduce redundancy, high-confidence contigs were dereplicated with dRep v3.4.0 [50], using a minimum contig length of 1,000 bp, a primary clustering threshold of 90% average nucleotide identity (ANI), and a secondary threshold of 95% ANI. The dereplicated set was re-analysed with Prodigal, and only genomes containing at least one complete open reading frame (partial=00 flag) were retained.

### Provisional taxonomic annotation and validation

Provisional annotations were obtained using BLASTx against the Virus-Host DB, with an e-value cutoff of 1 × 10⁻5, a minimum query coverage of 30%, and up to five hits per contig retained. These assignments were used to guide phylogenetic placement. To verify assembly quality, reads were mapped back to retained genomes with bwa-mem2 v2.2.1 [51] and coverage inspected in IGV v2.12.3 [52]. Terminal regions with inconsistent read support were trimmed prior to downstream analyses.

### Phylogenetic analysis

For tree reconstruction, only dereplicated contigs containing complete marker genes were used. The RNA-dependent RNA polymerase (RdRp) was selected for RNA viruses, and replication-associated proteins for DNA viruses. Open reading frames (ORFs) were predicted with NCBI ORFfinder [53]. Each marker ORF was compared to the NCBI nr database with BLASTp, and top hits were retrieved alongside representative sequences curated according to International Committee on Taxonomy of Viruses (ICTV) reference species.

Multiple sequence alignments were generated for each viral family or order using MAFFT v7.525 [54] with the --maxiterate 1000 --globalpair option to maximise alignment accuracy. Poorly aligned positions were removed with trimAl v1.5 [55], using a gap threshold of 0.75 and a block size of 10. Maximum-likelihood phylogenies were then reconstructed in IQ-TREE2 v2.3.4 [56], with branch support evaluated using 1,000 ultrafast bootstrap replicates. Resulting trees were rerooted manually in FigTree v1.4.4 [57] to optimise interpretability. Trees were visualised in RStudio v4.3.2 [58] using the ggtree package v3.17.1 [59].

### Viral abundance estimation and visualisation

Viral abundance was quantified following the approach of De Coninck et al. [23]. Reads were mapped back to the final dereplicated viral contigs using bwa-mem2 v2.2.1 [51], and CoverM v0.7.0 [60] was used to estimate abundance at the contig level. A contig was considered present within a pool if at least 50% of its length was covered by mapped reads. Read counts for viral contigs were summed per pool to generate an abundance matrix. This matrix was subsequently visualised in Rstudio using the pheatmap package [61].

### Ecological and geographic distribution of viral communities

To explore spatial and ecological patterns, viral presence–absence was determined at the site level based on contig detection criteria (≥50% breadth of coverage). Site-level matrices were collapsed across biological replicates and mapped to surveillance locations, with community composition visualised using piechart plots in R (scatterpie v0.2.1 [62]) overlaid on basemaps from rnaturalearth (v0.3.2 [63]). Putative arboviruses were defined as viruses assigned to genera containing recognised arboviral species, and were mapped separately to highlight their distribution across sites.

The relative abundance of viruses was calculated to assess variation across environmental and regional gradients. Viral read counts were aggregated at the family level, normalised within each site, and averaged across groups. Comparisons were made across land types (urban and rural) and first-level International Territorial Level (ITL1) regions of England and Wales. Visualisation was carried out using ggplot2 v3.5.1 [64].

To test for ecological associations, site-level presence–absence matrices were used to compare detection frequencies between urban and rural sites. Fisher’s exact tests were performed independently for each virus species and family, with false discovery rate (FDR) correction applied. In parallel, differential abundance testing was carried out at the site level to evaluate whether specific viral taxa were enriched in urban versus rural sites. Read counts were collapsed across replicates, aggregated by species or family, and analysed using DESeq2 (v1.36 [65]). To reduce the influence of rare taxa and spurious enrichment driven by highly skewed read distributions in a small number of samples [66,67], we applied a prevalence filter prior to differential abundance testing.

Specifically, we retained only families present in at least ∼20% of sites within both urban and rural groups (≥9 sites per group). Species-level patterns were examined only within families that passed this filter, to help identify potential contributors to family-level signals.

### Alpha and beta diversity analyses

To examine within- and between-site viral diversity, viral read counts were rarefied to a common depth corresponding to the 5th percentile of non-zero library sizes, with rarefaction repeated 1,000 times and mean diversity values retained.

Alpha diversity was quantified using observed richness (number of distinct viral taxa) and Shannon diversity, calculated in vegan v2.6-4 [68]. Diversity values were compared across land types (Rural and Urban) and mosquito species using Wilcoxon rank-sum tests, with p-values adjusted for multiple comparisons using the Benjamini–Hochberg procedure. Associations between alpha diversity and geographic coordinates (latitude and longitude) were first evaluated with Spearman rank correlations, and the strength of linear trends was subsequently assessed using least-squares regression.

Beta diversity was assessed using Bray–Curtis dissimilarities computed from relative abundance matrices. Ordinations were performed by principal coordinates analysis (PCoA) and non-metric multidimensional scaling (NMDS) in vegan, with ordination plots visualised in ggplot2. PERMANOVA (9,999 permutations) was used to test for effects of land type, latitude, and longitude on viral community composition, focusing on *Culex pipiens* to allow balanced comparisons across land types. Homogeneity of multivariate dispersion (PERMDISP) was evaluated using centroid-based distances.

An overview of the full experimental workflow is summarised in Figure 1.

**Figure 1.**
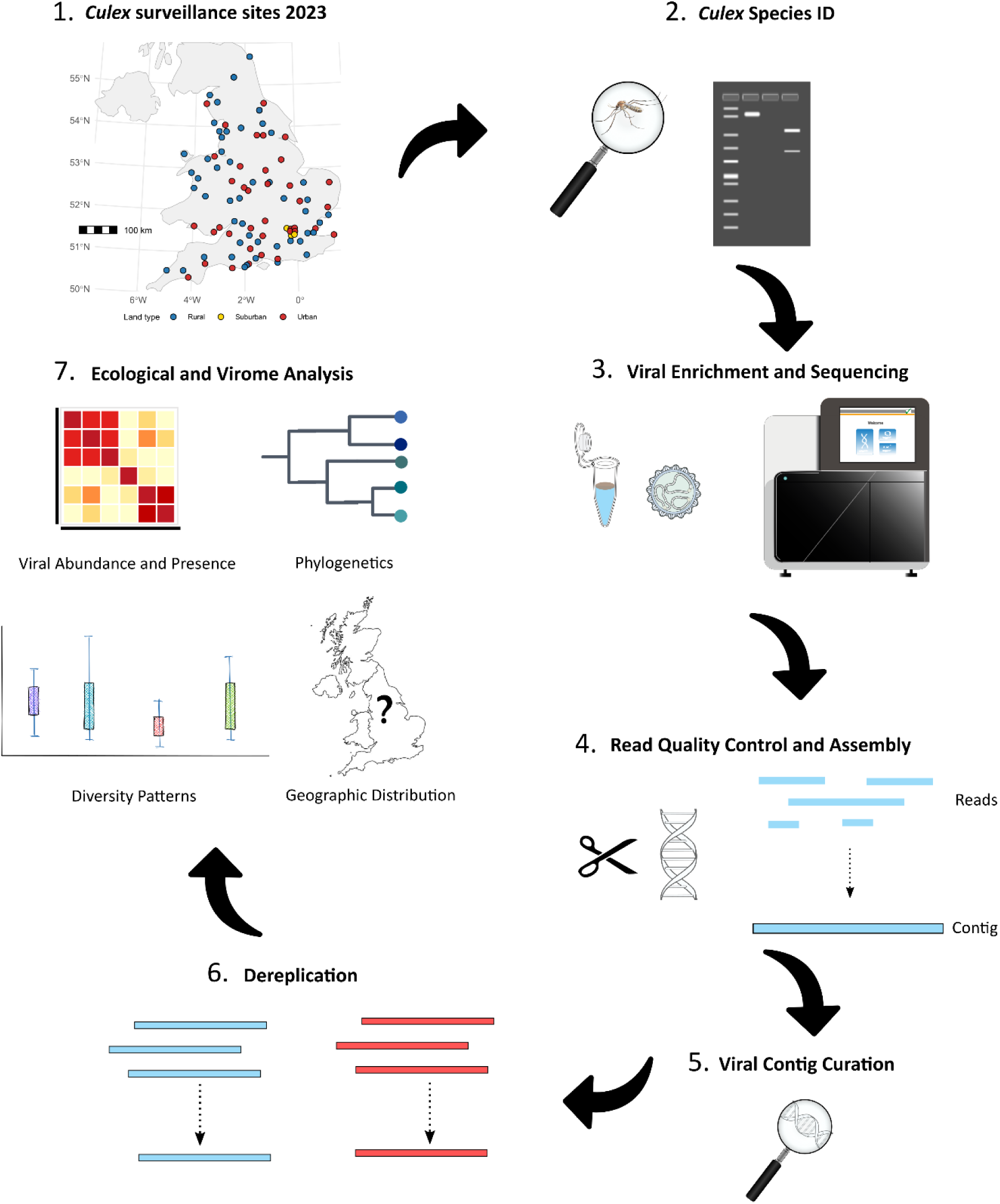
Schematic overview of the project workflow showing sampling, laboratory, and analytical steps used in the study.

## Results

### Taxonomic breadth and phylogenetic placement

The viral sequence processing pipeline yielded 253 dereplicated contigs containing at least one complete ORF across the 151 libraries. Among these, complete hallmark genes (RdRp for RNA viruses or Rep for DNA viruses) were recovered for 41 distinct taxa, spanning RNA viruses and a single DNA virus. At least one of these viruses was detected at 86 of the 93 sampled sites across England and Wales. These comprised negative-sense RNA viruses (n = 10), positive-sense RNA viruses (n = 22), double-stranded RNA viruses (n = 8), and a single-stranded DNA virus (n = 1). Phylogenetic reconstruction confirmed the placement of most lineages within recognised viral families, including *Iflaviridae* (n = 5), *Solemoviridae* (n=3), *Tymoviridae* (n = 3), *Peribunyaviridae* (n = 2), *Partitiviridae* (n = 2), *Rhabdoviridae* (n = 2), *Sedoreoviridae* (n = 2), *Orthomyxoviridae* (n = 2), *Xinmoviridae* (n = 2), *Amalgaviridae* (n = 1), *Chrysoviridae* (n = 1), *Chuviridae* (n = 1), *Dicistroviridae* (n = 1), *Draupnirviridae* (n = 1), *Mesoniviridae* (n=1), *Nodaviridae* (n=1). In addition, several sequences clustered outside established ICTV-designated viral families including 4 Negev-like viruses, Culex bunyavirus 2 (Order: *Hareavirales*), Daeseongdong-like virus 2, two *Ghabrivirales* spp., two *Tolivirales* spp. and one *Tymovirales* spp. (Table 1 and supplemental data).

**Table 1.** Summary of viruses detected in *Culex* spp., showing taxonomy based on phylogenetic placement and ICTV designation, mosquito hosts, nearest relatives, and whether novel or previously reported in the UK. ANI = Average Nucleotide Identity. Asterisks mark cases with no defined ICTV species threshold, where a 90% cutoff was applied based on the most commonly used standard.

| Virus name | Virus order | Virus family | Virus genus | Cx. pipiens | Cx. molestus | Cx. torrentium | ICTV Species demarcation (RDRP amino acid %) | Nearest hit accession (amino acid identity) | Novel virus | Previously detected in UK? |
| --- | --- | --- | --- | --- | --- | --- | --- | --- | --- | --- |
| Almendravirus Chester | Mononegavirales | Rhabdoviridae | Almendravirus | + | - | - | 90 | Almendravirus Chester; OZ253854.1 (100%) | N | Y |
| Alphamesonivirus fluvideense | Nidovirales | Mesoniviridae | Alphamesonivirus | + | + | + | 90* | Alphamesonivirus cavallyense; ALP32023.1 (100%) | N | Y |
| Amalgaviridae sp. | Durnavirales | Amalgaviridae | Unclassified | + | - | - | 90* | Tetrodontophora bielaniensis associated virus 1; DAB41738.1 (37%) | Y | N |
| Atherstone virus | Eliovirales | Peribunyaviridae | Orthobunyavirus | + | - | - | 96 | Atherstone virus; OZ253933.1 (100%) | N | Y |
| Chrysoviridae sp. | Ghabrivirales | Chrysoviridae | Alphachrysovirus | + | - | - | 70 | Chrysoviridae sp.; XLV26575.1 (100%) | N | N |
| Cripavirus pipiens | Picornavirales | Dicistroviridae | Cripavirus | + | - | - | 90 (capsid protein) | Aphis gossypii virus; UUG74202.1 (95%) | N | N |
| Culex bunyavirus 2 | Hareavirales | Unclassified | Unclassified | + | - | - | 90* | Culex bunyavirus 2; QRW41985.1 (100%) | N | Y |
| Culex circovirus-like virus | Jormunvirales | Draupnirviridae | Valentivirus | + | + | - | 80 ANI | Culex circovirus-like virus; OZ248129.1 (100%) | N | Y |
| Culex luteo-like virus | Sobelivirales | Solemoviridae | Unclassified | + | - | + | 90* | Culex luteo-like virus; WVU03164.1 (98%) | N | N |
| Culex mononega-like virus 1 | Mononegavirales | Xinmoviridae | Unclassified | + | + | - | 90* | Culex mononega-like virus 1; QGA70931.1 (100%) | N | N |
| Culex mononega-like virus 2 | Mononegavirales | Xinmoviridae | Unclassified | + | - | + | 90* | Guadeloupe mosquito mononega-like virus; QEM39177.1 (52%) | Y | N |
| Culex mosquito-like virus 4 | Jingchuvirales | Chuviridae | Culicidavirus | + | - | - | 90 | Culex mosquito virus 4; QRW42864.1 (99%) | N | Y |
| Culex Negev-like virus 1 | Unclassified | Unclassified | Unclassified | + | + | - | 90* | Culex Negev-like virus NS46; OZ251790.1 (99%) | N | Y |
| Culex Negev-like virus 2 | Unclassified | Unclassified | Unclassified | + | + | - | 90* | Negev virus; BAR91505.1 (100%) | N | N |
| Culex Negev-like virus 3 | Unclassified | Unclassified | Unclassified | + | - | - | 90* | Utsjoki negevirus 1; UYL94304.1 (31%) | Y | N |
| Culex Negev-like virus 4 | Unclassified | Unclassified | Unclassified | + | - | - | 90* | Culex Negev-like virus 1; UUG74013.1 (97%) | N | N |
| Culex pipiens betapartitivirus | Durnavirales | Partitiviridae | Betapartitivirus | + | - | - | 90 | Culex pipiens betapartitivirus 2; OZ253781.1 (100%) | N | Y |
| Culex pipiens deltapartitivirus | Durnavirales | Partitiviridae | Deltapartitivirus | + | - | - | 90 | Inari deltapartitivirus; UUV42371.1 (77%) | Y | N |
| Culex pipiens Ifla-like virus 1 | Picornavirales | Iflaviridae | Iflavirus | + | - | - | 90 (capsid protein) | Picornavirales sp.; QKN88975.1 (98%) | N | N |
| Culex pipiens Ifla-like virus 2 | Picornavirales | Iflaviridae | Iflavirus | + | - | - | 90 (capsid protein) | Culex Iflavi-like virus 4; YP_009552017.1 (98%) | N | N |
| Culex pipiens nodavirus | Nodamuvirales | Nodaviridae | Unclassified | + | - | - | 90* | Wufeng shrew nodavirus 5; WPV63049.1 (98%) | N | N |
| Culex pipiens Tymo-like virus 1 | Tymovirales | Tymoviridae | Unclassified | + | - | - | 90* | Nasturtium officinale macula-like virus 1; QQG34658.1 (62%) | Y | N |
| Culex pipiens Tymo-like virus 2 | Tymovirales | Tymoviridae | Unclassified | + | - | + | 90* | Lampyris noctiluca tymovirus-like virus 1; QBP37021.1 (59%) | Y | N |
| Culex pipiens Tymo-like virus 3 | Tymovirales | Tymoviridae | Unclassified | + | - | - | 90* | Sichuan mosquito tymo-like virus; UBJ25983.1 (91%) | N | N |
| Culex pipiens Tymovirales sp. | Tymovirales | Unclassified | Unclassified | + | - | - | 90* | Diaporthe helianthi tymovirus 1; WNM95042.1 (70%) | Y | N |
| Culex Sobemo-like virus | Sobelivirales | Solemoviridae | Unclassified | + | - | - | 90* | Plasmopara viticola lesion associated sobemo-like 1; QHD64767.1 (51%) | Y | N |
| Daeseongdong virus 2 | Unclassified | Unclassified | Unclassified | + | + | + | 90* | Orthornavirae sp.; XLV26731.1 (100%) | N | Y |
| Ghabrivirales sp. 1 | Ghabrivirales | Unclassified | Unclassified | + | - | + | 90* | Ghabrivirales sp.; XLV26802.1 (95%) | N | N |
| Ghabrivirales sp. 2 | Ghabrivirales | Unclassified | Unclassified | + | - | - | 90* | Ghabrivirales sp.; XLV26802.1 (100%) | N | N |
| Hedwig virus | Eliovirales | Peribunyaviridae | Gryffinivirus | + | + | - | 90 | Asum virus; QGA70944.1 (100%) | N | N |
| Ista virus | Picornavirales | Iflaviridae | Iflavirus | + | - | + | 90 (capsid protein) | Ista virus; OZ251434.1 (99%) | N | Y |
| Jotan virus | Picornavirales | Iflaviridae | Iflavirus | + | - | + | 90 (capsid protein) | Jotan virus; UYL94332.1 (99%) | N | N |
| Jotan-like virus | Picornavirales | Iflaviridae | Iflavirus | + | - | + | 90 (capsid protein) | Culex Iflavi-like virus 1; WVU03087.1 (84%) | Y | N |
| Marma virus | Sobelivirales | Solemoviridae | Unclassified | + | - | - | 90* | Marma virus; OZ251086 (100%) | N | Y |
| Merida virus | Mononegavirales | Rhabdoviridae | Merhavirus | + | - | - | 90 | Merida virus; AWJ96718.1 (97%) | N | N |
| Tolivirales sp. 1 | Tolivirales | Unclassified | Unclassified | + | - | - | 90* | Sanya tombus-like virus 2; UHM27571.1 (31%) | Y | N |
| Tolivirales sp. 2 | Tolivirales | Unclassified | Unclassified | + | - | - | 90* | Sanxia water strider virus 14; APG76440.1 (43%) | Y | N |
| Umatilla virus | Reovirales | Sedoreoviridae | Orbivirus | + | - | - | 78 | Umatilla virus; WZL41624.1 (99%) | N | N |
| Valmbacken virus | Reovirales | Sedoreoviridae | Unclassified | + | - | - | 90 | Valmbacken virus; WJJ55410.1 (100%) | N | N |
| Wuhan Mosquito virus 4 | Articulavirales | Orthomyxoviridae | Quaranjavirus | + | - | - | 90* | Wuhan Mosquito Virus 4; OZ251991.1 (100%) | N | Y |
| Wuhan Mosquito virus 6 | Articulavirales | Orthomyxoviridae | Quaranjavirus | + | + | - | 90* | Wuhan Mosquito Virus 6; AJG39092.1 (98%) | N | N |

In total, 11 viruses met ICTV criteria for novel species, with RNA-dependent RNA polymerase amino acid identities to their closest known relatives ranging from 31% to 84% (Table 1). All taxa were distinct based on dereplication and phylogenetic criteria, except *Ghabrivirales sp. 1* and *Ghabrivirales sp. 2*, which share 95 % amino-acid identity in the RdRp and are therefore considered a single provisional species under ICTV demarcation standards.

### Taxonomic highlights

Twelve viruses were detected in both this national survey and our previous zoo-based survey [37] (Table 1). The remaining detections represented taxa not previously observed in our earlier dataset. Among RNA viruses, members of the *Picornavirales* (five *Iflaviridae* and one *Dicistroviridae*) and *Mononegavirales* (four taxa) were prominent; phylogenetic analyses placed all of these within insect-specific clades (Supplemental data). Two taxa from the *Quaranjavirus* genus (Wuhan mosquito virus 4 and Wuhan mosquito virus 6) and four Negev-like viruses were identified, grouping with established mosquito-associated clades. Beyond insect-associated taxa, several lineages typically linked to plants or fungi were also present, including members of the *Solemoviridae*, *Chrysoviridae*, *Ghabrivirales*, *Partitiviridae*, and *Amalgaviridae*. One partitivirus matched Culex pipiens betapartitivirus 2, previously reported in the UK [37], while another represented a novel deltapartitivirus.

### Arbovirus-related detections

Beyond insect-specific lineages, several viruses closely related to recognised arboviruses were also detected (Figures 2 and 4B). Hedwig virus (Family: *Peribunyaviridae*; Genus: *Gryffinivirus*) was the most widespread, observed at 10 sites across southern England and Wales: Swansea, Newport, Bristol, Ipswich, Melton Mowbray, Newmarket, Slimbridge, Upper Stoke, and two London localities (Harlesden and Walworth). In six of these detections, complete RdRp ORFs were recovered, each showing >98% amino acid identity to previously reported Hedwig virus sequences (Figure 2B). Some clustered most closely with isolates from Germany, others with viruses reported from Sweden or France, indicating close relationships to multiple European lineages.

**Figure 2.**
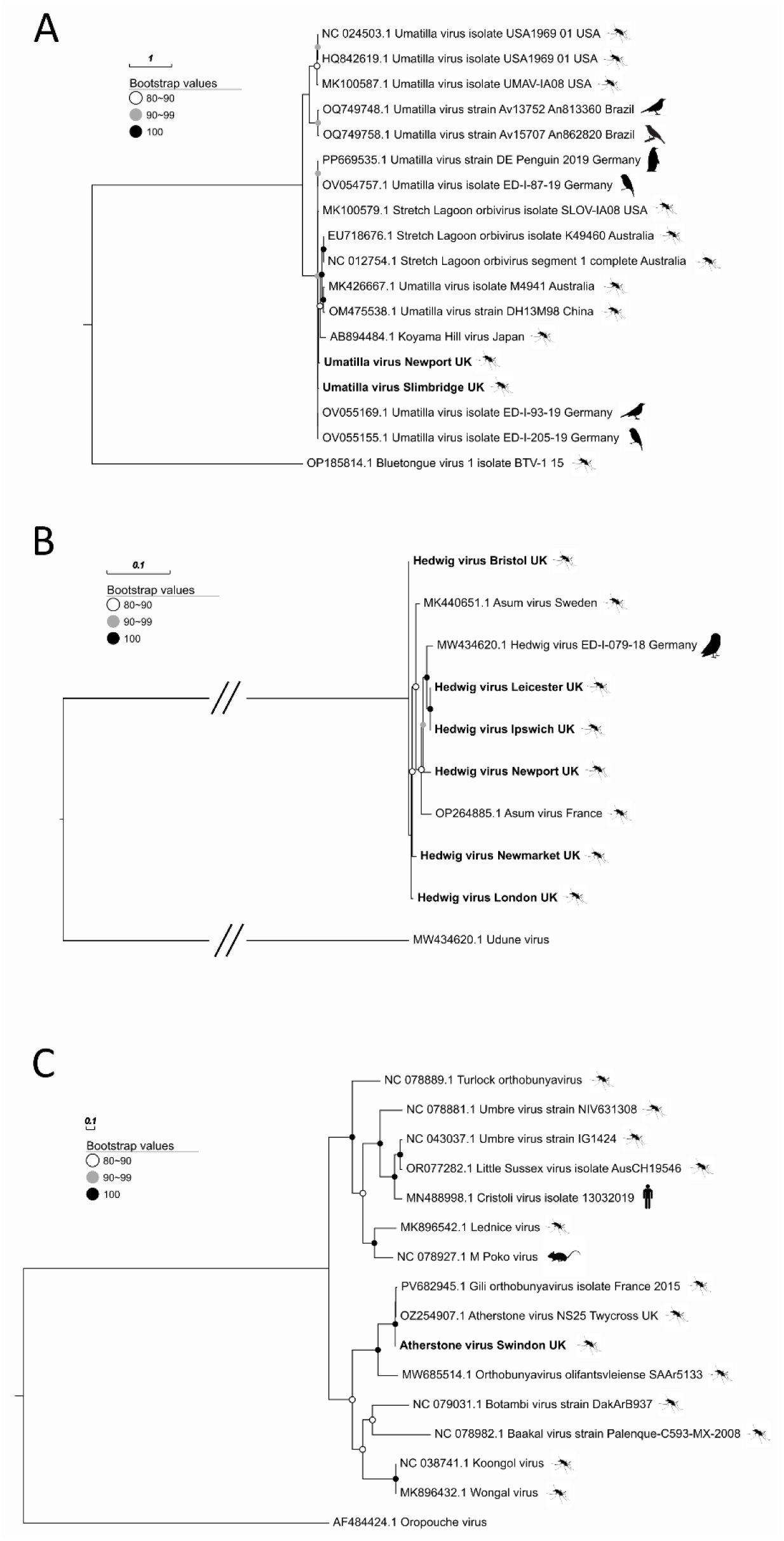
Maximum-likelihood trees of the RNA-dependent RNA polymerase (RdRp) ORFs of **(A)** Umatilla virus (*Sedoreoviridae; Orbivirus*) **(B)** Hedwig virus (*Peribunyaviridae; Gryffinivirus*) **(C)** Atherstone virus (*Peribunyaviridae; Orthobunyavirus*). Scale bars represent the number of amino acid substitutions per site. Silhouettes represent host source.

Umatilla virus (Family: *Reoviridae*; Genus: *Orbivirus*) was found at three sites, including Slimbridge, Newport and Plymouth. Representative sequences for the two sites clustered with others obtained from birds caught in Germany during 2019 surveillance (Figure 2A).

Atherstone virus (Family: *Peribunyaviridae*; Genus: *Orthobunyavirus*) was restricted to two sites in Swindon and Cambridge (Figure 2C and 4B), with RdRp genes showing near identical amino acid identity to the virus reported in our previous study [37] (Accession: OZ254907), and closely related to a partial sequence recently released from a detection in France from 2015 (Accession: PV682945).

For each of these viruses, all expected genome segments were recovered (except segment 3 of Umatilla virus) and co-occurred within single pools, confirming assembly of near-complete genomes rather than partial detections (ENA accessions: Hedwig virus – 3 segments [OZ335791-OZ335793]; Umatilla virus – 9 segments [OZ335966, OZ335967, OZ335972, OZ335978, OZ335979, OZ335986, OZ335988, OZ335991, OZ367119]; Atherstone virus – 3 segments [OZ335505, OZ335506, OZ335605]).

### Virus distribution patterns

Virus detections spanned a gradient from widespread to highly restricted taxa (Figure 3). Only three viruses, Daeseongdong virus 2, Wuhan mosquito virus 4, and Alphamesonivirus fluvideense, were widespread, each detected at more than a third of all sites and across all 10 ITL regions (Figures 3B and C). Sixteen viruses showed intermediate distributions, occurring at 6–25 sites, including *Chrysoviridae* sp. (25 sites), *Culex* Negev-like virus 1 (16 sites), and Marma virus (18 sites). By contrast, the majority of taxa (22/41) were restricted, being found at four or fewer sites, with eight observed only once. While most singletons have not previously been reported in *Culex* (e.g., *Amalgaviridae* sp., *Culex* Negev-like virus 3, *Culex pipiens* Tymo-like virus 1 and *Culex pipiens Tymovirales* sp.), others such as *Culex pipiens* betapartitivirus 2, Almendravirus Chester and Valmbacken virus have been documented in earlier studies [24,37,69], supporting their likely mosquito association despite low prevalence here.

**Figure 3.**
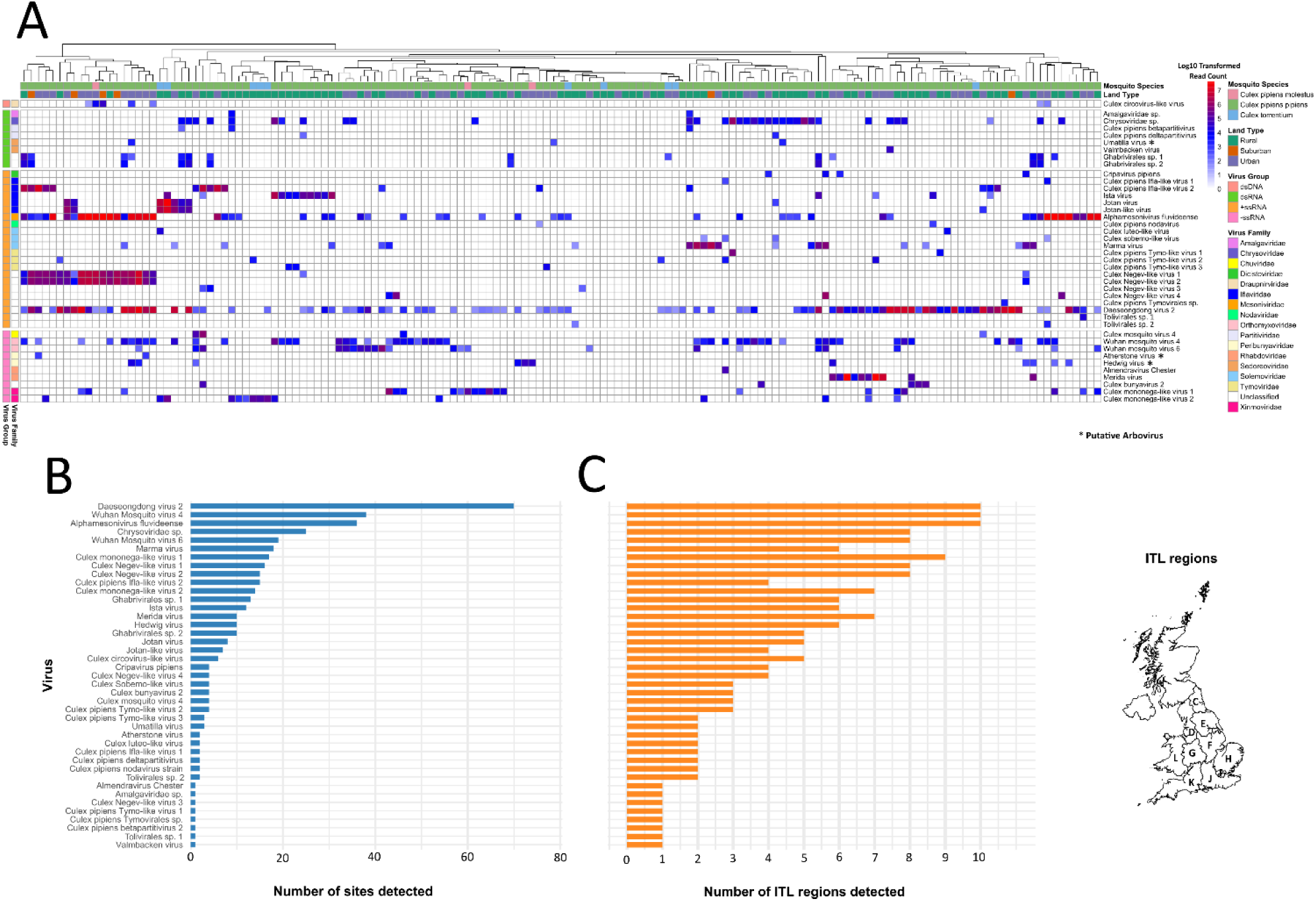
(A) Heatmap of virus reads detected in *Culex* spp. pools across 151 libraries from 93 sites. Number of sites **(B)** and ITL regions **(C**) each taxon was detected across.

Across 312 taxa detections spanning 86 sites, six families (*Mesoniviridae*, *Chrysoviridae*, *Orthomyxoviridae, Iffaviridae*, *Xinmoviridae*, and *Solemoviridae*) accounted for over half of all records (59%; Figure 4A). An additional 23% of detections fell into the ‘Unclassified’ category, reflecting viruses that could not be placed within established families. The majority of these undesignated detections reflected Daeseongdong virus 2, which was widespread, being detected at 70 sites across all 10 ITL regions (Figures 3B and C). Relative abundance profiles (read count) across ITL regions (Figure 4C), as well as land type (Figure 4D) were also dominated by these same families. At the detection (presence/absence) level, no viral families or species differed significantly in frequency between urban and rural sites (Fisher’s exact test, Table S1), suggesting no evidence of habitat-specific enrichment.

**Figure 4.**
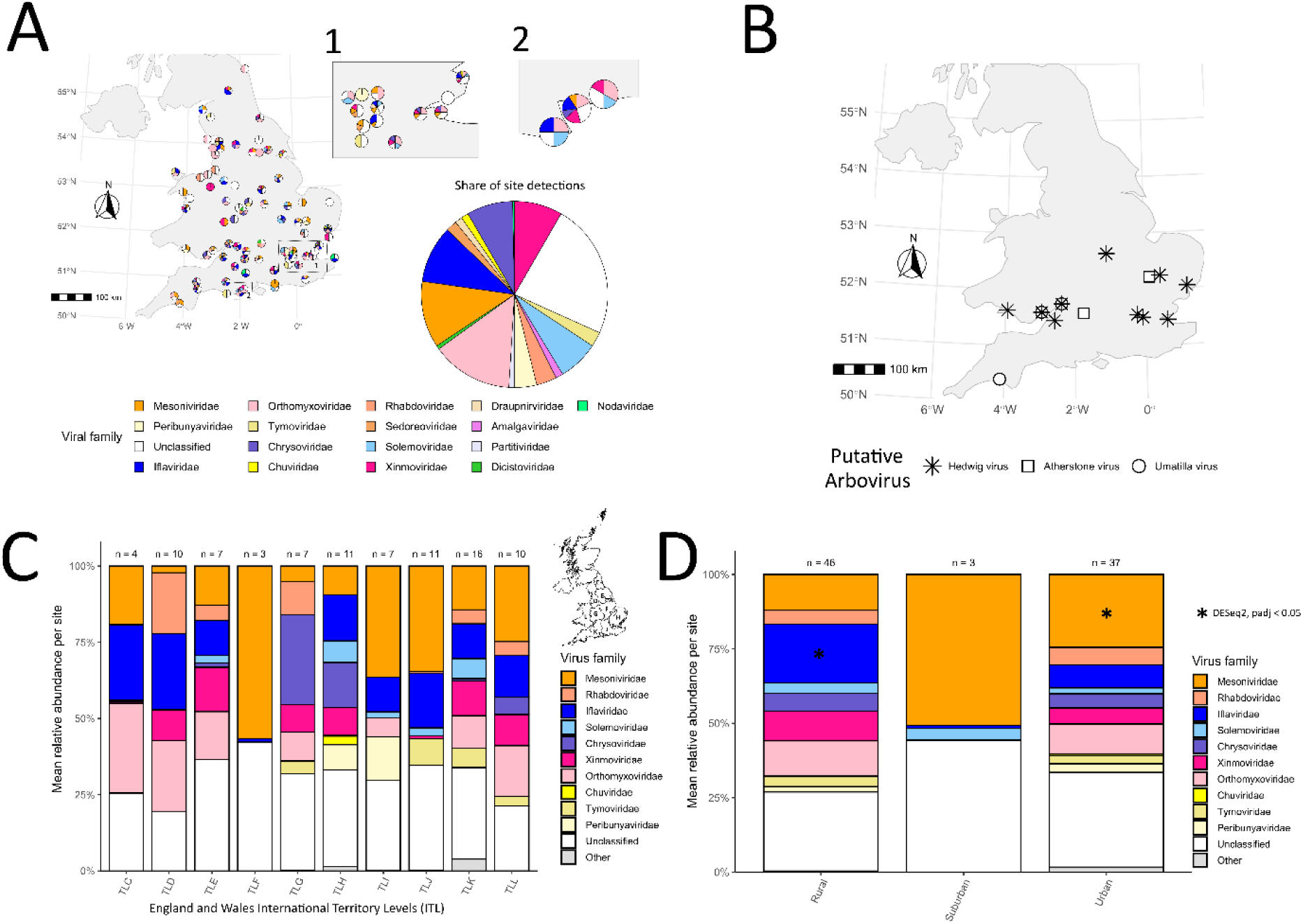
(A) Geographic distribution (presence/absence) of viral family detections across 86 of 93 sites where at least one taxon was detected. **(B)** Distribution of putative arboviruses across England and Wales. **(C)** Mean relative viral abundance (read counts) per site across the 10 ITL regions surveyed. **(D)** Mean relative viral abundance (read counts) per site by coarse land type. Asterisks denotes enriched taxa (rural vs urban).

In contrast, abundance-based comparisons (relative viral read counts) identified five families that met the prevalence filter for inclusion (>10% of both urban and rural sites): *Orthomyxoviridae*, *Mesoniviridae*, *Iflaviridae*, *Xinmoviridae*, and *Chrysoviridae*. Among these, only *Mesoniviridae* (urban-enriched, Deseq2 padj = 5.4 × 10⁻⁴) and *Xinmoviridae* (rural-enriched, Deseq2 padj = 0.031) showed significant differences.

### Diversity patterns

Viral richness and Shannon diversity did not differ significantly between urban and rural sites (Wilcoxon rank test, p > 0.05), with both measures showing similar ranges and dispersion within groups (Figures 5A–C). In *Cx. pipiens*, median richness was 2.6 (IQR 1.9) in rural sites and 2.5 (IQR 1.5) in urban sites, while Shannon diversity was likewise similar (rural: 0.34, IQR 0.65; urban: 0.32, IQR 0.62). For *Cx. torrentium* (n = 8) and *Cx. molestus* (n = 3), sample sizes were too limited for meaningful comparisons, though no clear land-use effect was evident.

**Figure 5.**
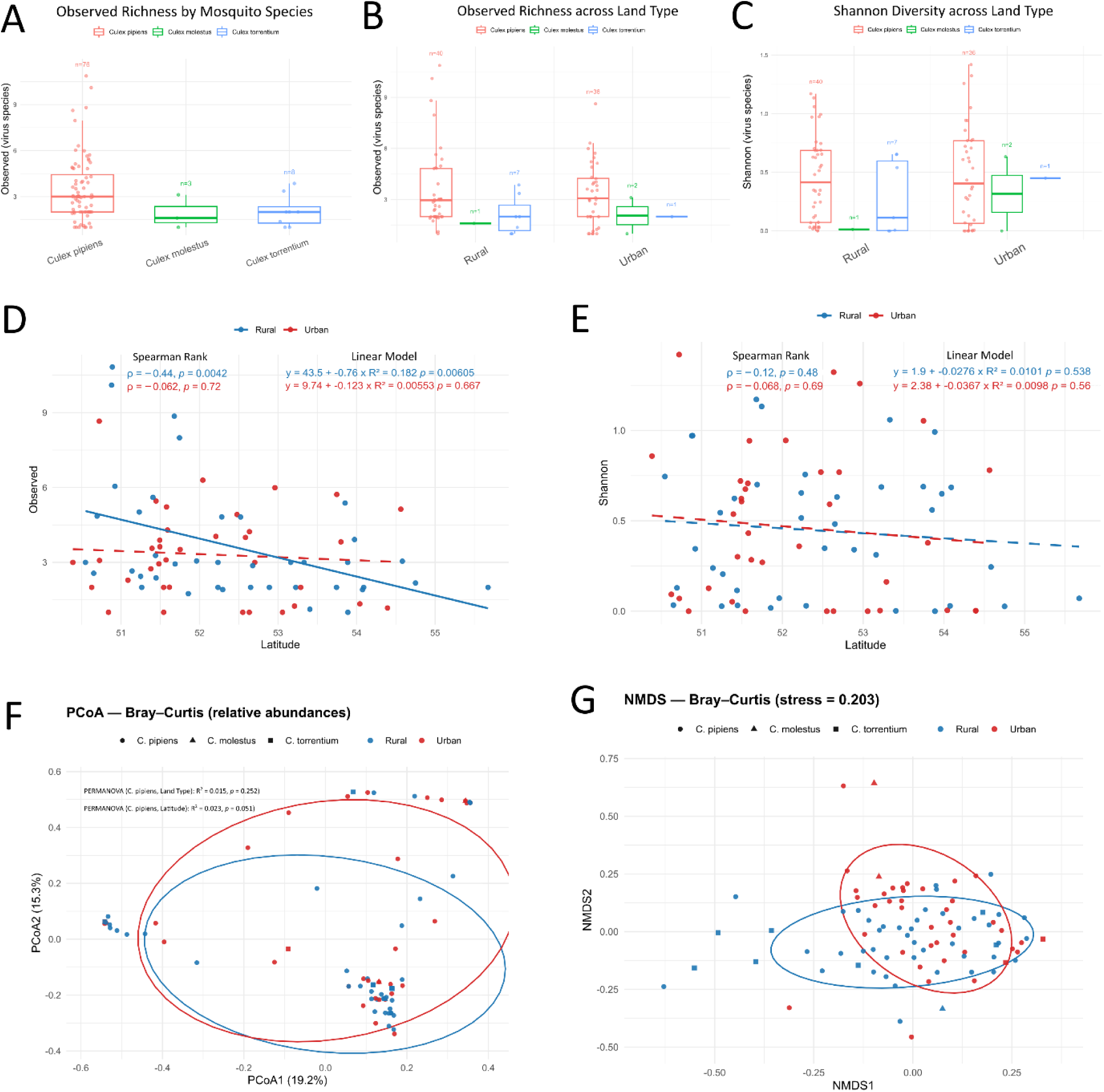
(A) Observed richness across *Culex* species **(B)** Observed richness stratified by species and compared between urban and rural sites **(C)** Shannon diversity stratified by species and compared between urban and rural sites **(D)** Association between latitude and observed richness in *Cx. pipiens pipiens* (solid line = statistical significance) **(E)** Association between latitude and Shannon diversity in *Cx. pipiens pipiens* **(F)** Principal coordinates analysis (PCoA) of Bray–Curtis dissimilarities in *Cx. pipiens pipiens*, with PERMANOVA testing effects of land type (urban vs. rural) and latitude. **(G)** Non-metric multidimensional scaling (NMDS) ordination of Bray–Curtis dissimilarities *in Cx. pipiens pipiens*.

Within *Cx. pipiens*, alpha diversity showed a significant negative association with latitude across rural sites for observed richness (Spearman’s ρ = –0.44, p = 0.0042; linear regression: R² = 0.182, p = 0.006), but not across urban sites (Spearman’s ρ = – 0.062, p = 0.72; linear regression: R² = 0.005, p = 0.667). This relationship was not detected for Shannon diversity, indicating that the number of viral taxa declined with increasing latitude but community evenness remained stable (Figures 5D and E). No associations with longitude were observed (See supplemental data).

Beta diversity analysis (Figure 5F) based on Bray–Curtis dissimilarities revealed no significant structuring of viral communities by land type (PERMANOVA: R² = 0.015, p = 0.264), but showed a borderline association with latitude (R² = 0.022, p = 0.051). Homogeneity of multivariate dispersion was confirmed (PERMDISP; rural mean 0.651 ± 0.011 SE, urban 0.630 ± 0.016 SE; p > 0.05), indicating that the lack of PERMANOVA significance reflected a true absence of compositional differences rather than unequal within-group variance. These results were consistent with the NMDS ordination, which showed broad overlap of communities across land types (Figure 5G).

## Discussion

This study represents the first comprehensive virome survey of UK *Culex* species, identifying 41 distinct viral taxa, among them three arbovirus-related lineages, including two detected for the first time in the UK. Our dataset, collected in the same year as the first UK WNV detection in *Aedes vexans* [12], found no evidence of WNV in *Culex* populations across any site, nor of USUV, the only mosquito-borne virus currently considered established in the UK [70]. By contrast, the detection of other putative arboviruses highlights potential emerging threats, spanning a continuum from neglected but increasingly reported (Umatilla virus) [71,72] to recently characterised with limited detection histories (Hedwig and Atherstone viruses) [37,72]. These results illustrate the added value of high-throughput sequencing as a complementary approach to existing UK surveillance, which has been primarily directed toward WNV and USUV.

Umatilla virus (UMAV), an orbivirus in the *Sedoreoviridae* family, was first isolated in the 1960s from *Culex* spp. collected in the USA [73]. It has since been detected in Australia [74], Japan [75], and Europe [71,72], and has re-emerged as a candidate pathogen in birds. In Germany, UMAV-positive blue tits were repeatedly reported with splenomegaly consistent with acute infection [72]. More strikingly, UMAV infection was confirmed in multiple deceased Cape penguins from a zoo, with one presenting with hepatitis and high viral loads across liver, spleen, and kidney [71]. In addition, a serological survey revealed high exposure rates in free-living pheasants, indicating frequent infection [71], suggesting that some avian species may serve as amplifying hosts, whereas others may be more prone to severe pathology.

Hedwig virus (HEDV), a species in the *Gryffinivirus* genus (*Peribunyaviridae*) was first reported in *Culex pipiens* from France in 2015, and has since been detected in mosquitoes in Sweden and Germany, as well as two birds (straw-necked ibis and ferruginous duck) [24,72]; of the necropsy reports available for these animals, pathological findings were inconsistent, leaving the pathogenic role of HEDV unresolved.

UMAV was detected at two sites near the Severn Estuary, while HEDV was detected at multiple sites near both the Severn and Thames Estuaries. This geographic overlap highlights estuarine regions as potential entry points for these viruses, consistent with proposed routes of historical incursions of vector-borne pathogens across Europe via migratory bird flyways [76–78].

Another member of the *Peribunyaviridae*, Atherstone virus, was first characterised in our previous zoo-based study [37]. Shortly after, it was reported from archived *Culex* pools in southern France, originally screened following the detection of Umbre virus in patient brain tissue [79]. Although Umbre virus was not identified in these mosquitoes, a partial sequence corresponding to Atherstone virus was recovered (reported as “Gili orthobunyavirus”). Together with additional detections at sites in Cambridgeshire and Wiltshire from the present study, these data indicate that Atherstone virus is more widely distributed than initially recognised. Unlike UMAV and HEDV, which already have vertebrate detections, Atherstone virus has so far been detected only in mosquitoes.

Nevertheless, its placement within the *Orthobunyavirus* genus, which includes multiple established arboviruses, and the presence of a vertebrate-specific virulence factor (non-structural S protein) [37,80] support its classification as a putative arbovirus and a priority candidate for studies on vector competence, host range, and potential health significance.

Beyond arboviruses, a further 38 distinct viral taxa were detected, with only three viruses detected at more than a third of sampling sites, indicating that most taxa were geographically restricted. A similar pattern was observed by Pan et al. [27], who reported that just 27 of 393 viruses were present in more than 25% of individual mosquitoes across China, likewise suggesting strong spatial structuring of mosquito viromes. In contrast, some studies, often based on large mosquito pools and short contigs annotated at broad taxonomic ranks (e.g. family level or by lowest common ancestor methods) rather than species-level phylogenetic classification, have reported a more conserved core virome [81,82]. Such higher-level analyses, however, tend to overestimate viral ubiquity by grouping genetically distinct species into broader taxonomic units, thereby masking underlying spatial and host-associated heterogeneity. Of the most prevalent taxa, Alphamesonivirus fluvideense was particularly notable. Although mesoniviruses are generally regarded as insect-specific [83], recent reports of Alphamesonivirus sequences in horse lung and lymph node tissues associated with respiratory disease [84] raise questions about their broader host associations. The significance of these vertebrate detections remains uncertain and may reflect rare spillover events, but the widespread occurrence of this lineage in this study highlights opportunities for exposure and underscores the value of including mesoniviruses in surveillance frameworks.

The interpretation of rarer viruses poses further challenges, as some may reflect dietary or environmental acquisition rather than true mosquito associations [85,86]. However, most singleton detections in this study involved viruses previously reported in *Culex* viromes (e.g., Valmbacken virus) or belonging to established mosquito-specific lineages (e.g., Negev-like viruses). Similarly, *Chrysoviridae*, *Solemoviridae* and *Partitiviridae* were once considered dietary contaminants but are now consistently recovered across independent *Culex* virome studies [23,24,81], indicating that they represent persistent mosquito-associated lineages.

The UK *Culex* virome showed little evidence of strong ecological structuring by coarse land type. Frequency-based comparisons indicated no habitat-specific enrichment of viral lineages, but abundance-based analyses suggested modest shifts, with *Mesoniviridae* more common in urban pools and *Xinmoviridae* more abundant in rural ones. These differences imply that while presence/absence patterns remain broadly consistent across habitats, virus abundances may still capture ecological contrasts such as variation in larval environments or feeding behaviours [87,88]. A modest but statistically significant negative correlation between latitude and viral richness in *Culex* pipiens across rural sites was also observed, with higher diversity observed in southern sampling locations. This pattern mirrors the broader latitudinal diversity gradients documented in viral and microbial communities [89,90]. These trends have been attributed to factors such as temperature-dependent insect activity and differences in environmental viral stability [91]. In the case of mosquitoes, warmer temperatures in southern regions may support longer activity periods and larger population sizes, thereby increasing opportunities for viral transmission and maintenance. The absence of this trend across our urban sites could reflect the urban heat island effect, which may buffer mosquitoes from broader latitudinal temperature differences [92,93].

The limited and inconsistent ecological structuring observed here highlights the challenges of incorporating virome data into arbovirus risk models. While metagenomic surveillance is clearly valuable for detecting circulating arboviruses, the heterogeneous distribution of ISVs, without evidence of a broad core virome or predictable structuring by land type, makes it difficult to parameterise their potential modulatory effects. Without clearer understanding of where and when particular ISVs are likely to occur, their influence on arbovirus transmission at a population level cannot be reliably incorporated into predictive frameworks.

Nonetheless, ISVs are increasingly considered as candidates for biological control [86]. For example, insect-specific flaviviruses have been shown to inhibit the replication of arbo-flaviviruses such as WNV and Dengue fever viruses [32,33]. However, the absence of Insect-specific flaviviruses in our dataset is consistent with other European *Culex* virome investigations [23,24,69,81].

In contrast, we detected several other ISVs of potential interest, including *Bunyaviricetes* members such as Culex bunyavirus 2, which has been reported previously from *Culex* populations [28,37,94], as well as four *Tymovirales* species (*Alsuviricetes*), two of which are novel. Given their phylogenetic proximity to arboviruses of concern in Europe, such as Rift Valley fever virus (*Bunyaviricetes*), as well as chikungunya and Sindbis viruses (*Alsuviricetes*), these lineages represent logical candidates for targeted evaluation. Rift Valley fever virus has not shown local transmission in continental Europe but remains a priority for surveillance and preparedness due to the risk of introduction and establishment through animal movements [95]. In contrast, chikungunya virus has already caused repeated autochthonous outbreaks in southern Europe [14,96], while Sindbis virus is endemic in parts of northern Europe, where it occasionally causes human infections [13].

## Conclusions

As arbovirus threats increase in temperate regions, agnostic virus screening approaches should be seen as essential components of vector-borne disease preparedness. This study exemplifies this with the detection of two known arboviruses previously undetected in the UK and a further putative arbovirus with unknown health impacts. Beyond arboviruses, the UK Culex virome appears heterogeneous, with limited evidence of a conserved core or ecological structuring, aside from a modest latitudinal gradient in richness across rural sites. This suggests that viral communities are shaped predominantly by fine-scale or stochastic processes. Building on this national baseline, longitudinal sampling will be essential to capture temporal dynamics and evaluate broader virome stability. In parallel, integration of accumulating virome data with vertebrate surveillance, including serology in birds, mammals and humans, will help clarify host associations and refine arbovirus risk evaluation.

## Data availability

Virus sequences and raw reads are available at the European Nucleotide Archive under project accession number PRJEB98260.

## Author contributions

MB, MSCB and JM secured funding for this project. JP, MSCB and ACD contributed to the conceptual development of the project. EW, RW, AGCV, JM, MB and MSCB co-ordinated fieldwork. JP and EW conducted laboratory work. JP carried out bioinformatic analysis. JP, MSCB and ACD interpreted data. JP produced figures and drafts of the manuscript. All authors assisted in critical revision of the manuscript.

## Acknowledgements

We thank Agata Delnicka, Amelia Simpson, Anthony J. Abbott, Colin J. Johnston, Jude Martin, Kendall Barlow, Eloise Aliski, Saffron Shiels, Sara Gandy, and Sarah M. Biddlecombe for their invaluable assistance with mosquito field collections. We also acknowledge the support of Richard Gregory and the Centre for Genomic Research (CGR) at the University of Liverpool for providing access to computational resources used in this research, as well as for the use of CGR’s sequencing facilities.

