## Supplementary figures and images for "Mosquito Viromes in England and Wales Reveal Hidden Arbovirus Signals and Limited Ecological Structuring"

### supplemental data

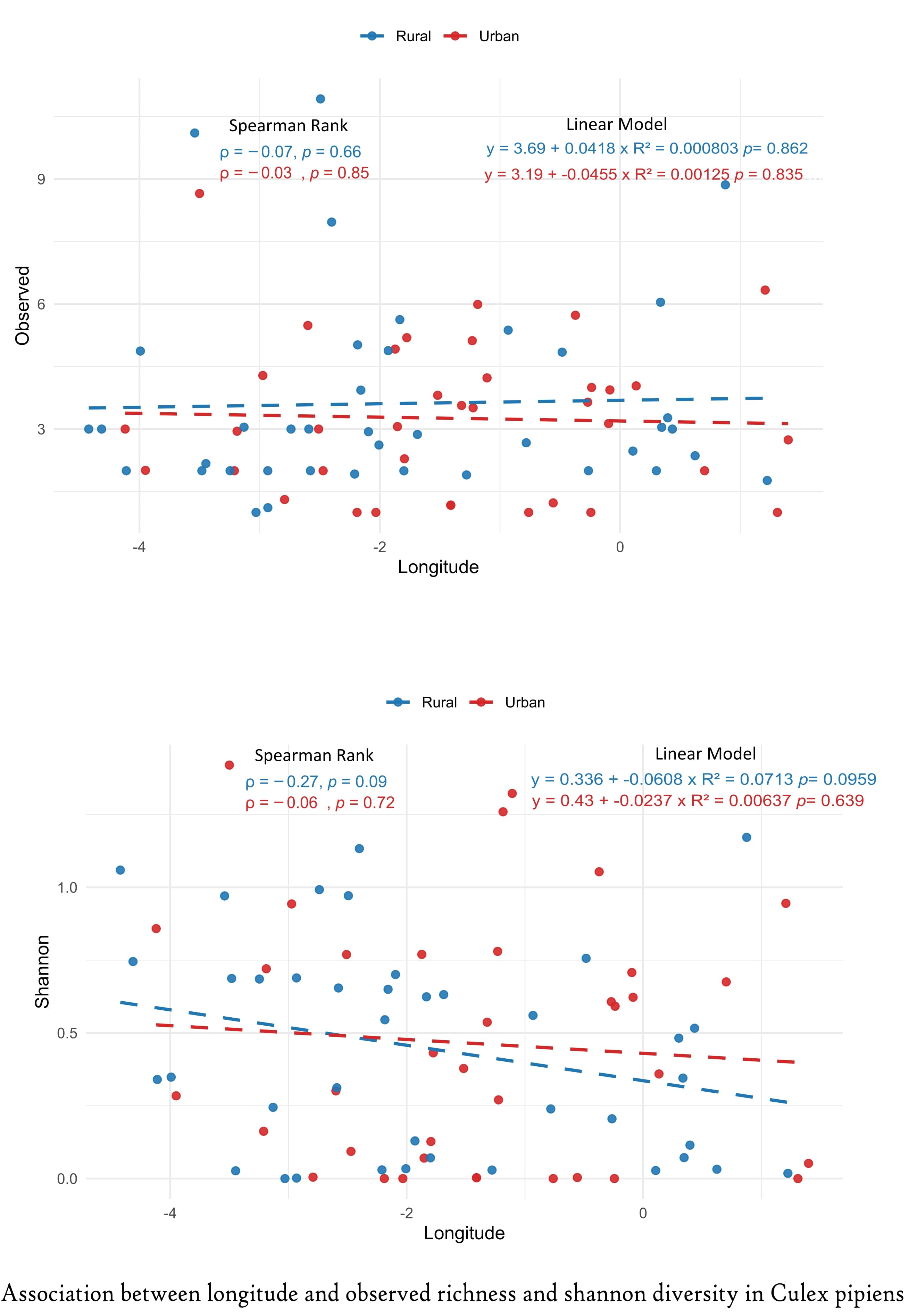

### supplemental data

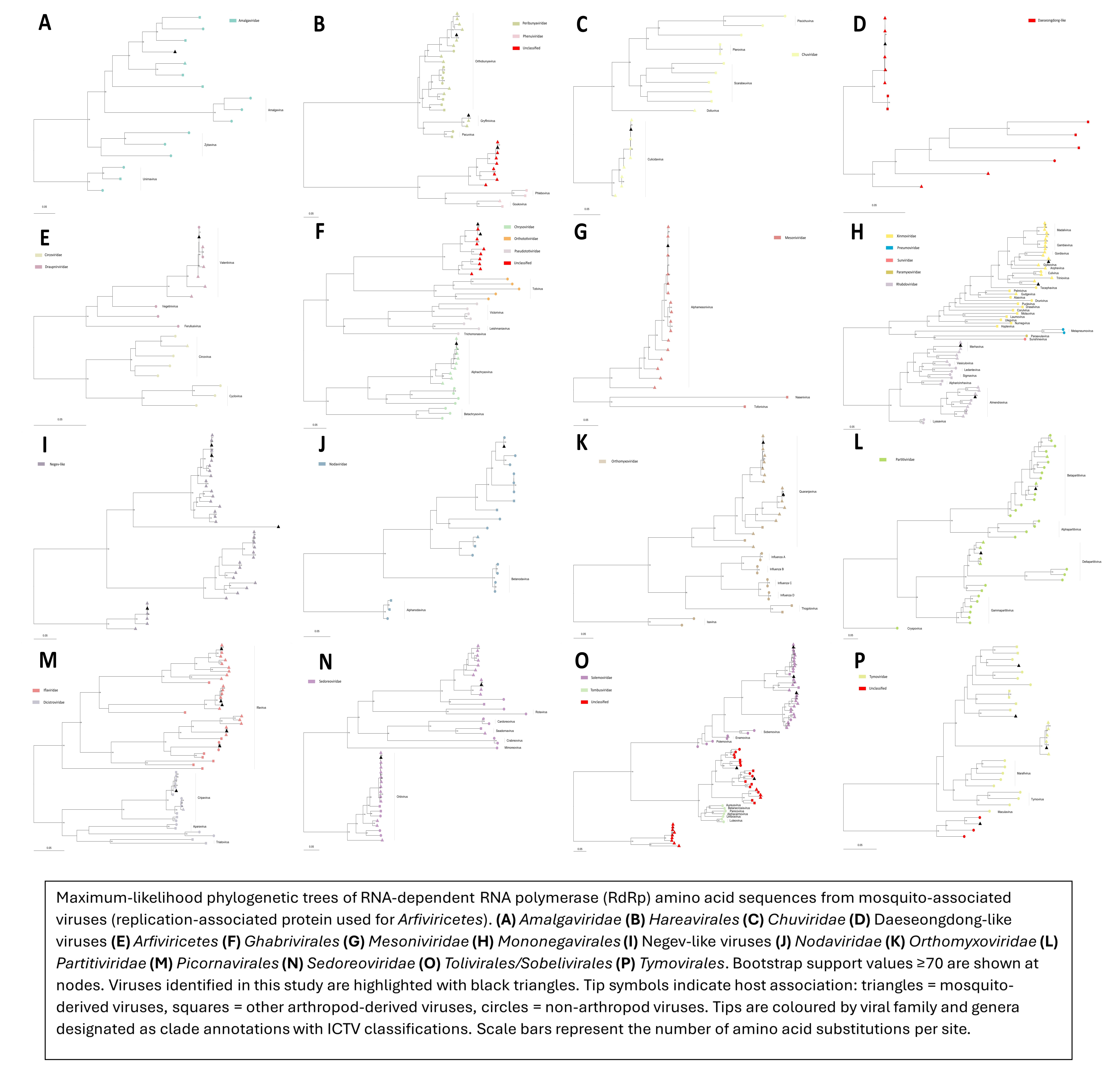
